# The genomic signature of social interactions regulating honey bee caste development

**DOI:** 10.1101/012385

**Authors:** Svjetlana Vojvodic, Brian R. Johnson, Brock Harpur, Clement Kent, Amro Zayed, Kirk E. Anderson, Timothy A. Linksvayer

**Affiliations:** Department of Entomology, University of Arizona, USA; USDA Carl Hayden Bee Research Center, USA; Department of Entomology, University of California Davis, USA; Department of Biology, York University, Canada; Department of Biology, University of Pennsylvania, USA

**Keywords:** indirect genetic effects, interacting phenotypes, extended phenotype, social evolution

## Abstract

Social evolution theory posits the existence of genes expressed in one individual that affect the traits and fitness of social partners. The archetypal example of reproductive altruism, honey bee reproductive caste, involves strict social regulation of larval caste fate by care-giving nurses. However, the contribution of nurse-expressed genes, which are prime socially-acting candidate genes, to the caste developmental program and to caste evolution remains mostly unknown. We experimentally induced new queen production by removing the current colony queen, and we used RNA sequencing to study the gene expression profiles of both developing larvae and their care-giving nurses before and after queen removal. By comparing the gene expression profiles between both queen-destined larvae and their nurses to worker-destined larvae and their nurses in queen-present and queen-absent conditions, we identified larval and nurse genes associated with larval caste development and with queen presence. Of 950 differentially-expressed genes associated with larval caste development, 82% were expressed in larvae and 18% were expressed in nurses. Behavioral and physiological evidence suggests that nurses may specialize in the short term feeding queen- versus worker-destined larvae. Estimated selection coefficients indicated that both nurse and larval genes associated with caste are rapidly evolving, especially those genes associated with worker development. Of the 1863 differentially-expressed genes associated with queen presence, 90% were expressed in nurses. Altogether, our results suggest that socially-acting genes play important roles in both the expression and evolution of socially-influenced traits like caste.

## Introduction

The social insect sterile worker caste is the archetypal example of reproductive altruism that initially puzzled Darwin(Darwin 1859) and spurred Hamilton(Hamilton 1964) to develop kin selection theory. Kin selection theory presupposes the existence of genes that are expressed in one individual but have fitness effects on relatives(Hamilton 1964). Despite this clear focus of social evolution theory on socially-acting genes, empirical studies of the genetic basis of social insect traits, including caste, have widely overlooked the potential effects of genes that are expressed in one individual but affect the traits of social partners.

Honey bee female caste is considered to be an exemplar polyphenism, whereby the expression of alternate queen and worker morphs is controlled by environmental cues(Evans and Wheeler 1999). Unlike some other well-studied polyphenisms that are controlled by simple abiotic factors such as temperature or photoperiod(Nijhout 2003), honey bee queen-worker dimorphism critically depends on social control of larval development by adult nestmates(Linksvayer et al. 2011). *In vitro* rearing studies demonstrate that in the absence of social control, queen-worker dimorphism disappears and a continuous range of phenotypes are produced(Linksvayer et al. 2011).

Honey bee colonies only rear new queens during specific life history stages, for example in the spring when the colony is large enough to split in half, or upon the death of the current queen. Queen rearing is an emergent, colony-level process involving the coordinated activities of hundreds or thousands of adult workers. Necessary steps include the construction of special queen cells by nurse bees (Fig. 1), distinct provisioning behavior of nurses coupled with distinct qualitative and quantitative differences in the nutrition fed to queen- and worker-destined larvae (colloquially known as “royal jelly” vs. “worker jelly”)(Haydak 1970; Brouwers et al. 1987), the larval developmental response to these environmental signals, and finally, selection by nurses of a subset of larvae in queen cells to be reared to adulthood(Hatch et al. 1999).

**Figure 1.**
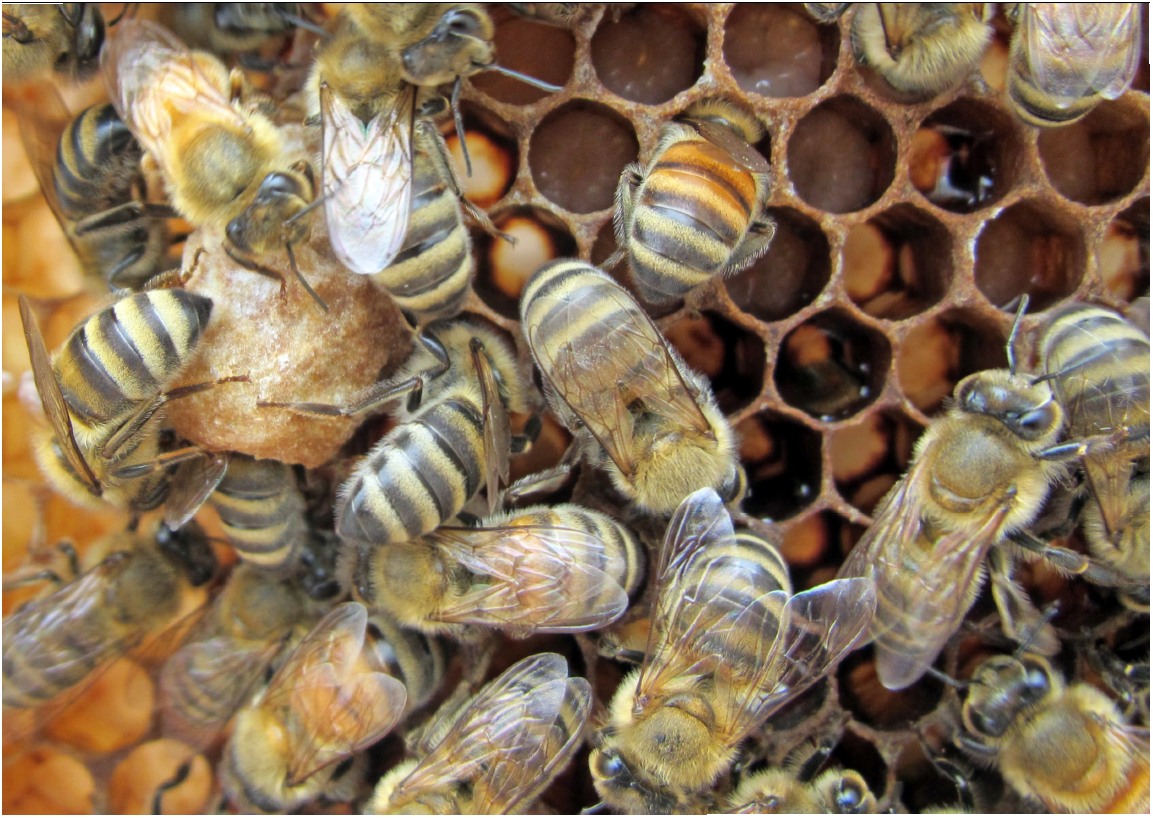
Honeybee workers rear most of their larvae in hexagonal cells (upper right) provisioned with a relatively small quantity of food so that the larvae develop into new workers. A few larvae are reared as new queens in larger queen cells (center left) that are newly constructed and provisioned with more and qualitatively different brood food.

Previous studies of the genetic basis of caste and other social insect traits have mainly used a conventional genetic approach, which seeks direct links between an individual’s genotype or patterns of gene expression and its phenotype(Evans and Wheeler 1999; Barchuk et al. 2007; Chandrasekaran et al. 2011). These studies have led to exciting progress in our understanding of the endogenous molecular genetic, epigenetic, and endocrine basis of alternate larval developmental trajectories in response to socially-controlled nutritional inputs(Evans and Wheeler 1999; Barchuk et al. 2007; Kucharski et al. 2008; Foret et al. 2012). For example, experimental gene knockdown studies demonstrate that insulin/TOR pathways mediating physiological and developmental responses to the nutritional environment strongly affect an individual’s caste fate(Patel et al. 2007; Mutti et al. 2011; Wolschin et al. 2011). However, the conventional approach has limited ability to identify exogenous socially-acting genes(Hahn and Schanz 1996; Wolf and Moore 2010).

As a result, the contribution of genes expressed in adult nestmates (e.g., nurses and foragers) to the genetic basis and evolution of the honey bee caste developmental program has received relatively little attention. Two exogenous, nurse-produced royal jelly proteins have been implicated as promoting queen development(Kamakura 2011; Huang et al. 2012). These and other protein-coding genes are very highly expressed in nurse worker hypopharyngeal and mandibular glands(Santos et al. 2005; Jasper et al. 2014), and different proportions of these glandular secretions are combined with sugars and proteins and fed to larvae, depending on the age and caste trajectory of the larva(Haydak 1970; Brouwers et al. 1987). Social control of caste development means that exogenous molecular factors expressed in adult nestmates may make up a significant portion of the colony-level gene regulatory network underlying queen development(Linksvayer et al. 2011). Indeed, quantitative genetic studies have demonstrated that the expression of honey bee caste and caste-related traits depends on both larval genotype and nurse genotype(Osborne and Oldroyd 1999; Beekman et al. 2000; Linksvayer et al. 2009a; Linksvayer et al. 2009b).

We use a novel extension of the conventional genetic approach to begin to characterize the full set of molecular interactions underlying social interactions that regulate reproductive caste development(Linksvayer et al. 2012), or the “social interactome” of caste. Instead of searching only for associations between an individual’s own patterns of gene expression and its traits, we also search for associations between social partners’ patterns of gene expression and the traits of focal individuals. Specifically, we used RNA sequencing of queen- and worker-destined larvae as well as “royal nurses” and “worker nurses”, nurses collected in the act of feeding queen- and worker-destined larvae, respectively. Subsequently, we used a new honey bee population genomic dataset(Harpur et al. 2014) to study how rates of molecular evolution vary at the genes we identified as being associated with the caste developmental program. We also determined whether there was evidence for behavioral and physiological specialization of nurses to feed queen- versus worker-destined larvae, as such specialization is expected to strengthen the transcriptional signature of social effects on caste development.

## Methods

### Basic setup, honey bee behavioral observations, and sampling

We conducted two studies of queen and worker rearing at the USDA Carl Hayden Bee Research Center in Tucson, AZ in April and June 2011, using commercial *Apis mellifera* stock colonies to create 4-frame observation hives. The first study focused on behavioral observations of individually marked workers that were involved in rearing new queens and workers, and the second focused on collecting nurse and larval samples for RNA sequencing. Observation hives were constructed with a hinged plexiglass door over each frame on each side so that it was possible to gently open the door and collect nurse and larval samples without disturbing the colony. The studies mimicked emergency queen rearing that occurs in the days immediately following queen loss.

The first study used two replicate observation hives. Every three days beginning 24 days before the start of the study, we individually marked 400 newly emerged adult workers with a unique combination of numbered tag glued onto the mesosoma and an age-specific abdomen paint mark, and we added 200 individually marked workers to each observation hive. Frames of known-aged brood were produced by caging queens on empty frames for 24 hours and then checking for the presence of eggs. Four days later, one frame with only similarly-aged 1^st^ instar larvae was placed into each observation hive, and the queen was removed to initiate emergency queen rearing. These frames were the source of young focal larvae, a fraction of which were reared as new queens, and the rest as workers. Within the first two days of queen removal, nurse workers build wax queen cells over young focal brood and begin provisioning these queen-destined larvae differentially than worker-destined larvae in worker cells (Figure 1). We continually observed areas of the frame with focal brood that contained both queen cells and worker cells and recorded the date, time, and identity of nurses observed provisioning queen or worker cells (i.e. “royal nurses” or “worker nurses”). Feeding behavior was defined when workers had their head positioned deep enough into the worker or queen cell to be in contact with the larva, and remained motionless except for a rhythmic motion of the abdomen for at least 5 seconds.

The second study used three replicate observation hives. The setup followed the first study, except that we collected samples of focal brood under both queen-present and queen-removed conditions. First, on the fourth day after introducing focal brood, samples of five 4^th^ instar worker-destined larvae and 20 nurses observed feeding 4^th^ instar worker-destined larvae were collected. Two days later, a new frame of same-aged 1^st^ instar larvae was added to each of the three observation hives, and each colony queen was removed in order to initiate emergency queen rearing. On the fourth day after introducing focal brood and removing the queen, we collected five 4^th^ instar worker-destined larvae from the frame of focal brood, and we collected 20 worker nurses in the act of feeding these 4^th^ instar worker focal brood. Similarly, we collected 20 royal nurses in the act of provisioning 4-day-old queen cells. Finally, we collected five 4^th^ instar queen larvae from the 4 day old queen cells. After removal from the hive, samples were immediately frozen in liquid nitrogen and stored on dry ice. We chose to collect larval and nurse samples when the larvae were 4^th^ instar because this is a period of very rapid larval growth(Haydak 1970; Evans and Wheeler 1999; Barchuk et al. 2007) as well as when differences in nurse provisioning are marked(Haydak 1970), even though most caste-related characters are considered to be already determined by this stage(Dedej et al. 1998).

In total, we collected: (1) worker larvae colonies with a queen, (2) worker larvae from queenless colonies, (3) queen larvae from queenless colonies, (4) worker nurses from colonies with a queen, (5) worker nurses from queenless colonies, and (6) royal nurses from queenless colonies. Thus, for both larvae and nurses, there were three conditions that were associated with: the production of new workers in hives with a queen; the production of new workers in queenless hives; and the production of new queens in queenless hives. We extracted RNA from whole larvae, but from nurses we dissected the two main glandular sources of proteinaceous brood food, the hypopharyangeal glands (HPG) and mandibular glands (MG), as well as the remaining head tissue (H, including brains and salivary glands).

### Nurse tissue dissections and mRNA sequencing

Nurse heads were thawed in RNA*later* (Qiagen), immediately dissected, and the three tissues (HPG, MG, H) collected and stored in RNA*later* at -80. In order to quantify HPG gland size variation within and between conditions, we took an image at 50x of a small subsample of each HPG, and three haphazardly-chosen HPG acini were measured at their widest point by an observer blind to the sample treatment.

RNA was extracted from individual larval samples and from tissue pooled from 5 nurses, for each of the three nurse tissue types, using Qiagen RNeasy kits. RNA concentration was quantified with Nanodrop and final pools created by combining RNA from 5 larvae from each of the 3 replicate colonies (15 total larvae), or from a tissue from 20 nurses from each of the 3 replicate colonies (60 total nurses). Separate pools were created for each of the three conditions and four tissues (L, HPG, MG, H), resulting in 12 total pools. RNA sequencing libraries were constructed at the University of Arizona Genetics Core, using RNA TruSeq library construction kits and Bioanalyzer RNAchips to check the library quality prior to sequencing. RNA samples were multiplexed on an Illumina HiSeq2000 with 6 samples per lane on two lanes with 100bp paired-end reads. Sequences were post-processed through trimmomatic to remove Illumina adapter sequences. Fastx and cutadapt software packages were used to remove reads with average quality scores < 25 and the ends of reads were clipped so that the mean quality of the last five bases was > 25. To control for initial variation in raw read number among samples within tissues, we used a standardized number of raw reads across all samples within each tissue to control for initial variation. Raw data summarizing the reads mapped to each gene and total number of reads uniquely aligned and unambiguously mapped for all samples will be deposited at DataDryad. Raw read data will be deposited at the NCBI SRA archive.

### Differential gene expression analysis

We aligned the reads to the *Apis mellifera* genome build 4.5(Elsik et al. 2014) using Tophat v2.04(Trapnell et al. 2012) with Bowtie2 and default parameters. We used htseq-count in the HTSeq(Anders and Huber 2010) Python Package with default parameters to assemble transcripts and obtain RPKM (Reads Per Kilobase per Million mapped reads) counts, based on the *A. mellifera* Official Gene Set 3.2(Elsik et al. 2014). We subsequently used two different R v3.1.0 (www.r-project.org) packages to analyze differential gene expression, EBSeq v1.5.4(Leng et al. 2013) and DEseq2 v1.4.5(Love et al. 2014). EBSeq uses an empirical Bayesian approach to identify the most likely among multiple possible expression patterns. We considered three alternatives: 1. the null hypothesis that no samples had differential expression; 2. the alternative hypothesis that expression in the sample associated with queen development/rearing was different than the samples associated with worker development/rearing; and 3. the alternative hypothesis that expression in the sample with the queen present was different than expression in the samples with the queen removed. We used default settings except for an increased number of iterations (maxround=40) to ensure convergence. With DESeq2 we used default settings and ran two separate analyses to identify genes with differential expression associated with queen- vs. worker-development and genes with differential expression associated with queen presence vs. absence. We focus on the EBSeq results for subsequent analyses because EBSeq is most appropriate for our study, but we also report DESeq2 results because the DEseq2 analysis was more conservative for identifying genes associated with caste (Fig. S1), but not for genes associated with queen presence (Fig. S2). Subsequent analyses were qualitatively similar following either EBSeq and DESeq2 differential expression analysis. Finally, we annotated transcripts with Blast2go(Conesa et al. 2005) and performed Gene Ontology (GO) enrichment analysis with the GOstats(Falcon and Gentleman 2007) R package. Venn Diagrams of differentially-expressed genes were constructed with the VennDiagram(Chen and Boutros 2011) R package.

### Molecular evolution analysis

To study patterns of molecular evolution at our identified differentially-expressed nurse and larval genes, we compared the estimated strength of selection on the genes since the divergence of *A. mellifera* and *A. cerana*, ^∼^5-25 Mya(Harpur et al. 2014). Specifically, we used a new database of estimates of the population size-scaled selection coefficient *γ* (*γ* = 2*N*_*e*_*s*; the product of effective population size and the average selection coefficient)(Harpur et al. 2014). These estimates are based on a population genomic comparison of polymorphism at synonymous and nonsynonymous sites within an African *A. mellifera* population compared to fixed differences between *A. mellifera* and *A. cerana*(Harpur et al. 2014). We compared *γ* estimates for differentially-expressed genes to background genes, which were not differentially expressed but had expression levels summed across all samples that were greater than or equal to the minimum expression levels in the list of differentially-expressed genes. Finally, we compared *γ* estimates for different categories of caste-associated genes.

## Results

### Analysis of nurse behavioral and physiological specialization

To clarify the potential specialization of nurses on provisioning worker vs. queen cells, we observed the feeding behavior of individually marked workers in two colonies over a period of 4 days during emergency queen rearing, for a total of 40 hours of observation. Nurses observed provisioning queen cells were on average 1.6 days younger than nurses observed provisioning worker cells (9.3 vs. 10.9 days, respectively; Fig. 2) (glm, quasipoisson residuals, *t* = 2.60, *df* = 191, *P* = 0.01). Of individual nurses observed for multiple feeding events within a single day, 37 provisioned only queen cells or worker cells, and 13 provisioned both. Of those observed multiple times among days, 7 provisioned only queen cells or worker cells and 11 provisioned both. Thus, nurses tended to provision only queen cells or worker cells within days but not across days (Fisher’s Exact Test, *P* < 0.001). We also measured the size of nurse HPG acini as an indicator of gland activity(Ohashi et al. 2000). Using residuals after controlling for differences among replicate colonies, royal nurses had larger HPG acini than worker nurses in queen present conditions (Tukey contrast with glm, *z* = 2.94, *P* = 0.009), but all other comparisons were not different (Tukey contrasts with glm, all *P* > 0.19) (Fig. 3).

**Figure 2.**
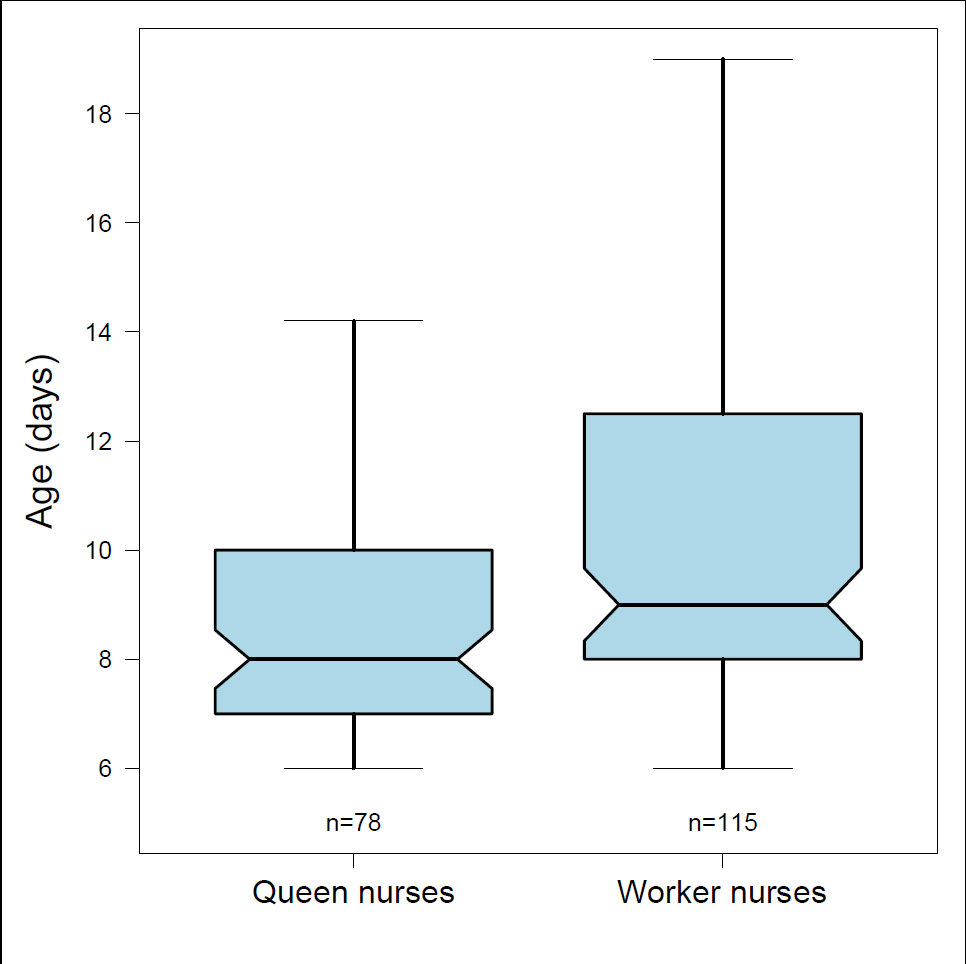
Box and whisker plot of the age of individually-marked “royal nurses” that were observed feeding queen-destined larvae in queen cells compared to “worker nurses” that were observed feeding worker-destined larvae in worker cells. Outliers are removed for clarity.

**Figure 3.**
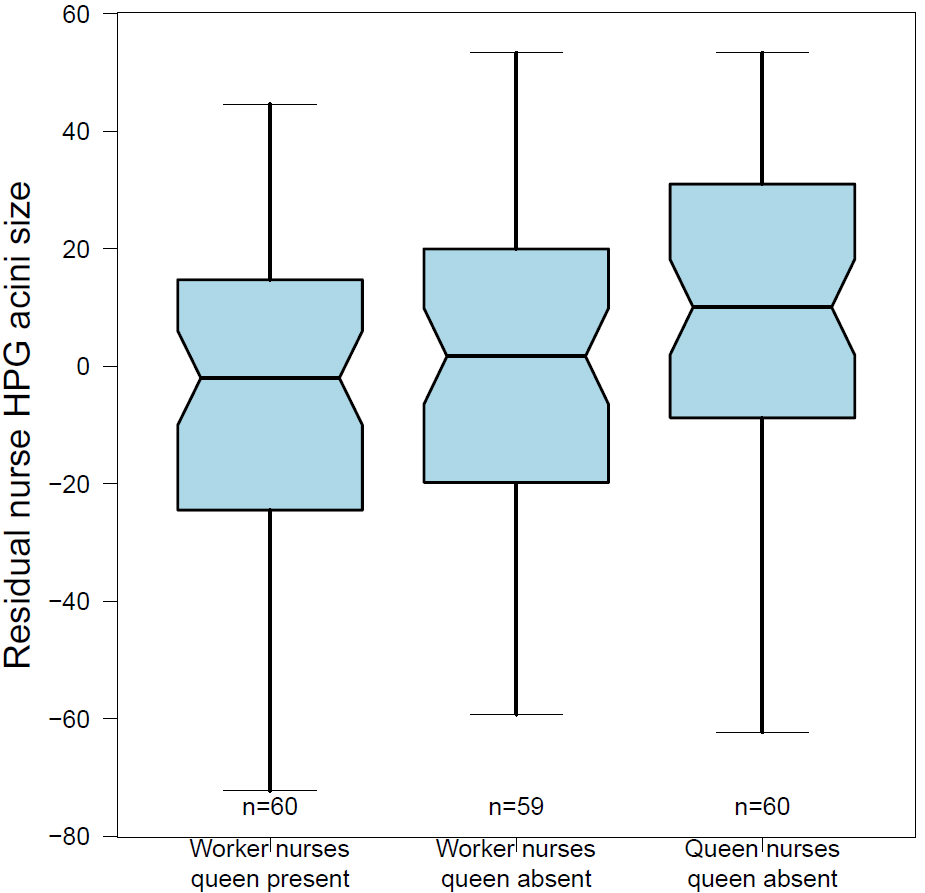
Box and whisker plot of residual nurse hypopharyngeal gland acini size (μm) depending on queen presence and nurse provisioning behavior. Royal nurses had larger HPG acini than worker nurses collected from colonies with a queen. Outliers are removed for clarity.

### Differential expression analysis

We identified 950 differentially expressed genes associated with whether larvae developed into new queens or workers (Table 1; Table S2). As expected, the majority of these genes (82%; 779/950) were differentially expressed in the larvae themselves, depending on whether the larvae were queen- or worker-destined larvae. 18% (171/950) were differentially expressed in *nurses* collected while feeding queen-destined larvae compared to nurses collected while feeding worker-destined larvae (3 expressed in MG, 105 H, and 63 HPG) (Table S2). Overlap of differentially-expressed genes associated with caste development is shown by tissue type in Figure S3.

**Table 1.** Select differentially-expressed nurse genes associated with caste development. Mean expression across conditions shown as LoglOFPKM, relative expression in royal nurse tissues vs. worker nurse tissues shown as Log2FoldChange, tissue (H = head tissue, MG = mandibular gland tissue), whether the gene was upregulated in royal nurses or worker nurses, annotation, inferred functional category, and whether the encoded protein has been identified in the royal jelly proteome, and thus assumed to be secreted from nurse glands to the brood food.

| Gene | Log10FPKM | Log2FoldChange | Tissue | Up-regulated | Annotation | Function | RJ proteome |
| --- | --- | --- | --- | --- | --- | --- | --- |
| GB53576 | 5.80 | 1.36 | H | royal | apisimin precursor | antimicrobial | yes |
| GB53576 | 5.80 | -1.39 | MG | worker | apisimin precursor | antimicrobial | yes |
| GB41428 | 4.10 | 1.65 | HPG | royal | defensin-1 preproprotein | antimicrobial | yes |
| GB51223 | 2.81 | 1.91 | HPG | royal | hymenoptaecin preproprotein | antimicrobial | yes |
| GB51223 | 2.51 | 2.52 | H | royal | hymenoptaecin preproprotein | antimicrobial | yes |
| GB47318 | 1.71 | 1.52 | HPG | royal | abaecin precursor | antimicrobial |  |
| GB53578 | 3.98 | 1.18 | H | royal | glucosylceramidase-like isoform 1 | metabolic activity | yes |
| GB43805 | 2.93 | 0.82 | H | royal | membrane metallo-endopeptidase-like 1-like | metabolic activity | yes |
| GB55204 | 5.58 | 0.88 | H | royal | major royal jelly protein 3 | nutritional | yes |
| GB45796 | 5.38 | 1.07 | H | royal | major royal jelly protein 3- partial | nutritional | yes |
| GB50012 | 3.73 | 0.99 | HPG | royal | hypothetical protein LOC726323 | unknown | yes |
| GB50012 | 3.36 | 1.51 | H | royal | hypothetical protein LOC726323 | unknown | yes |
| GB49583 | 2.36 | 1.50 | HPG | royal | 40s ribosomal protein s14 | protein synthesis |  |
| GB50709 | 2.00 | 1.22 | HPG | royal | 40s ribosomal protein s19a-like | protein synthesis |  |
| GB45374 | 2.99 | 0.66 | HPG | royal | 40s ribosomal protein s23-like | protein synthesis |  |
| GB50356 | 3.42 | 1.58 | HPG | royal | 60s acidic ribosomal protein p2-like | protein synthesis |  |
| GB52789 | 2.61 | 1.80 | HPG | royal | 60s ribosomal protein l22 isoform 1 | protein synthesis |  |

We also identified 2069 genes that were differentially expressed depending on queen presence, i.e. whether the mother queen was present or removed, irrespective of larval caste fate or nurse behavior (Table 2; Table S3). 90% (1863/2069) were expressed in nurse tissues, especially MG (1744 MG, 105 H, 15 HPG), and 206 were expressed in larval tissue. Overlap of differentially-expressed genes associated with queen presence is shown by tissue type in Figure S4.

Considering the top 25 most highly expressed genes for each tissue (Table S1), 40% (10/25) were shared among the nurse tissues. Many of these highly-expressed nurse genes are known to have protein products that are present in royal jelly(Schonleben et al. 2007; Furusawa et al. 2008; Zhang et al. 2014) (Table S1). Approximately one third of each set of most highly-expressed genes was unique to each nurse tissue, whereas ^∼^90% (22/25) of the most highly expressed larval genes were unique to larvae (Fig. S5).

**Table 2.** Select differentially-expressed nurse genes associated with caste development. Mean expression across conditions shown as LoglOFPKM, relative expression in royal nurse tissues vs. worker nurse tissues shown as Log2FoldChange, tissue (H = head tissue, MG = mandibular gland tissue), whether the gene was upregulated in royal nurses or worker nurses, annotation, inferred functional category, and whether the encoded protein has been identified in the royal jelly proteome, and thus assumed to be secreted from nurse glands to the brood food.

| Gene | Log10FPKM | Log2FoldChange | Tissue | Up-regulated | Annotation | Function | RJ proteome |
| --- | --- | --- | --- | --- | --- | --- | --- |
| GB55205 | 5.42 | 0.85 | H | queen present | major royal jelly protein 1 precursor | nutrition | yes |
| GB55212 | 4.70 | 1.21 | H | queen present | major royal jelly protein 2 precursor | nutrition | yes |
| GB55211 | 3.94 | 0.84 | H | queen present | major royal jelly protein 2 precursor | nutrition | yes |
| GB55206 | 4.03 | 0.75 | H | queen present | major royal jelly protein 4 precursor | nutrition | yes |
| GB55208 | 3.99 | 0.79 | H | queen present | major royal jelly protein 5 | nutrition | yes |
| GB55209 | 5.17 | 0.84 | H | queen present | major royal jelly protein 5 precursor | nutrition | yes |
| GB55207 | 3.21 | 0.86 | H | queen present | major royal jelly protein 6 precursor | nutrition | yes |
| GB55213 | 4.10 | 0.66 | H | queen present | major royal jelly protein 7 precursor | nutrition | yes |
| GB55215 | 2.14 | 1.44 | H | queen present | major royal jelly protein 9 precursor | nutrition | yes |
| GB55729 | 2.89 | -1.03 | MG | queen absent | major royal jelly protein 1 | nutrition | yes |
| GB45797 | 2.39 | 1.79 | MG | queen present | major royal jelly protein 1- partial | nutrition | yes |
| GB55205 | 5.72 | -1.39 | MG | queen absent | major royal jelly protein 1 precursor | nutrition | yes |
| GB45796 | 5.39 | 0.77 | MG | queen present | major royal jelly protein 3- partial | nutrition | yes |
| GB55208 | 4.25 | 1.93 | MG | queen present | major royal jelly protein 5 | nutrition | yes |
| GB55209 | 5.28 | 0.79 | MG | queen present | major royal jelly protein 5 precursor | nutrition | yes |
| GB55207 | 3.28 | -0.48 | MG | queen absent | major royal jelly protein 6 precursor | nutrition | yes |
| GB55213 | 4.39 | -0.25 | MG | queen absent | major royal jelly protein 7 precursor | nutrition | yes |

GO enrichment analysis for differentially-expressed genes associated with caste or queen presence are shown by tissue type in Tables S4 and S5, respectively. Among genes associated with caste: genes differentially expressed in nurse HPG tissue were enriched for GO terms associated with translation and several categories associated with immune function; nurse head tissue genes showed a weaker signal of enrichment for a range of GO terms, including signaling; and larval-expressed genes were enriched for terms such as metabolic processes and chromatin assembly. Among genes that were differentially expressed depending on queen presence: nurse MG genes were enriched for a range of terms including translation and transcription, macromolecular biosynthesis, signal transduction, metabolism, and immune response; nurse head tissue genes were enriched for immune system function, brain development, and chromatin assembly; and larval genes were enriched for terms such as response to oxidative stress and metabolism.

### Molecular evolution analysis

Differentially-expressed genes, whether associated with caste development or queen presence had higher average selection coefficients (γ) than non-differentially expressed genes (Fig. 4; glm on log-transformed gamma estimates, all *P* < 10^−8^), and furthermore, genes with expression associated with caste or both caste and queen presence had higher *γ* than genes with expression only associated with queen presence (Fig. 4; Tukey contrasts, both *P* < 10^−4^).

**Figure 4.**
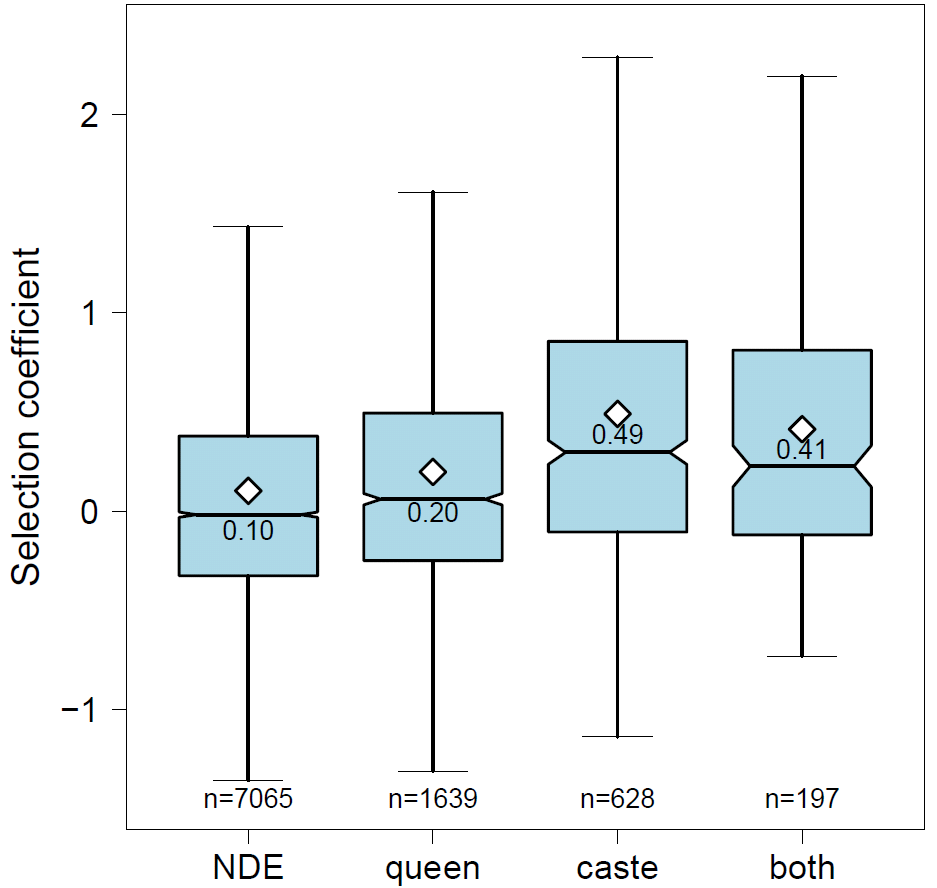
Box and whisker plot of population size-calibrated selection coefficients (γ) for non-differentially expressed genes (NDE), nurse and larval genes with expression associated with queen presence (”queen”), nurse and larval genes with expression associated with caste development (”caste”), and nurse and larval genes with expression associated with both queen presence and caste in different tissues (”both”). Means are indicated by white diamonds and also printed in each box. Outliers are removed for clarity.

Next we focused on genes with caste-associated expression. To further compare patterns of molecular evolution at genes associated with queen vs. worker production, we defined genes up-regulated in queen larvae or royal nurse tissues as “queen-associated genes” and genes up-regulated in worker larvae or worker nurse tissues as “worker-associated genes”. Mean *γ* for worker-associated genes was higher than queen-associated genes (glm with log-link on *γ* + 2 values, *t* = 2.47, *df* = 824, *P* = 0.014), and did not depend on whether the genes were expressed in larval or nurse tissues (*P* = 0.33) (Fig. 5). When only considering nurse-expressed genes, *γ* was higher for queen-associated vs. worker-associated genes (*t* = 3.71, *df* = 135, *P* = 0.0076), but *γ* was not significantly different when only considering larval-expressed genes (*t* = 1.78, *df* = 688, *P* = 0.076).

**Figure 5.**
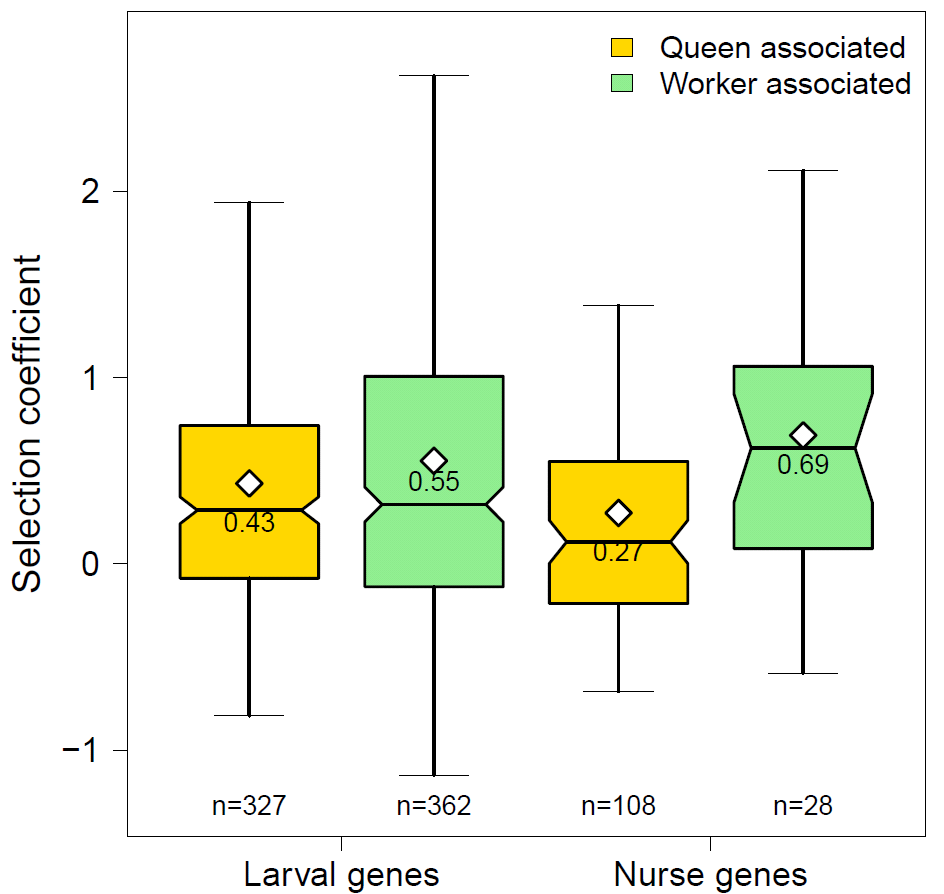
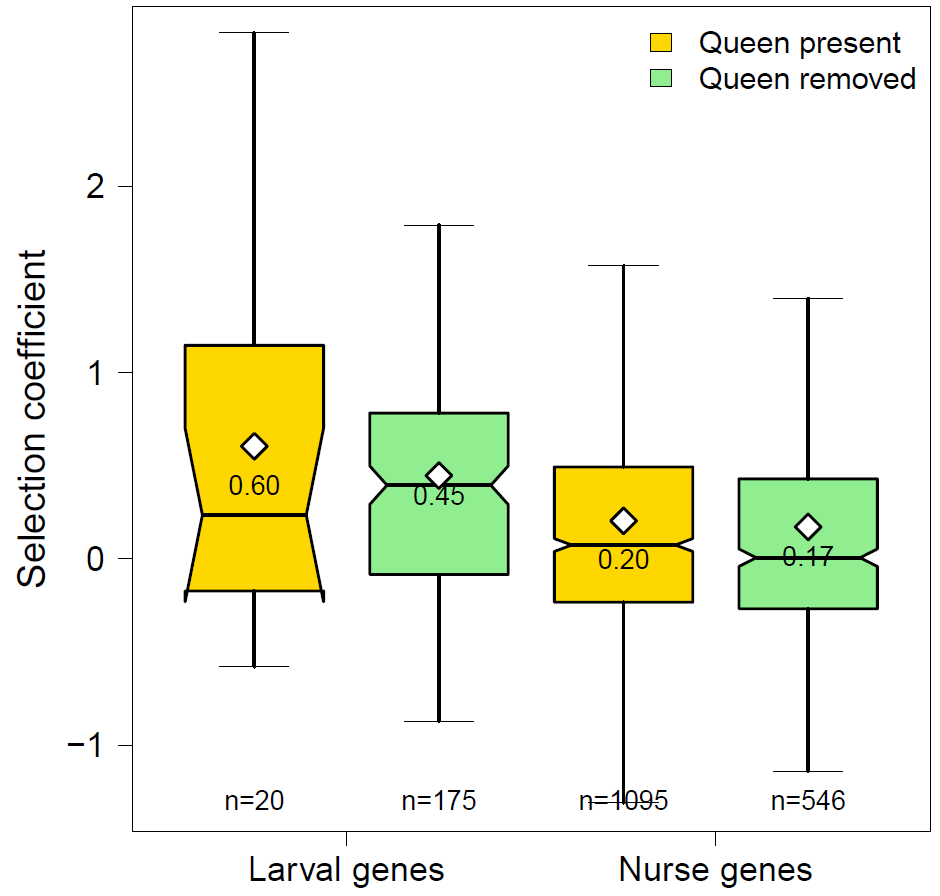
Box and whisker plot of population size-calibrated selection coefficients (γ) for nurse and larval differentially-expressed genes associated with caste. Genes are grouped by tissue type (larval vs. nurse tissues), and whether they were up-regulated in queen larvae or royal nurses (queen associated, yellow boxes) or they were up-regulated in worker larvae or worker nurses (worker associated, green boxes). Means are indicated by white diamonds and also printed in each box. Outliers are removed for clarity.

## Discussion

We simultaneously studied the gene expression profiles of two classes of socially-interacting individuals – developing larvae and their care-giving nurses – in order to identify genes expressed in larvae and their nurses that are associated with larval caste development. Such an approach has recently been used to study the molecular basis of host-parasite interactions(Tiemey et al. 2012; Westermann et al. 2012), but until now had only been proposed as a means to study the molecular basis of social interactions(Linksvayer et al. 2012). Increasingly, studies have shown how the gene expression profiles of many animals, including honey bees, ants, fruit flies, and cichlid fish strongly depend on the social environment(Grozinger et al. 2003; Robinson et al. 2008; Malka et al. 2014; Manfredini et al. 2014). Social environments in turn depend on the traits – and genes – of social partners(Wolf and Moore 2010). With such interdependence, the simultaneous study of the traits and genes of interacting partners is likely needed to capture the full dynamic social interplay affecting behavior, physiology, development, trait expression, and fitness(Johnson and Linksvayer 2010; Linksvayer et al. 2012).

We identified hundreds of genes that were differentially expressed in both developing honey bee larvae and care-giving nurse workers that were associated with whether the larvae were destined to develop as new queens or workers. The majority of these genes (82%; 779/950) were differentially expressed in the larvae themselves, depending on larval caste trajectory. These larval-expressed genes are assumed to be directly involved in the expression of developmental plasticity underlying queen-worker dimorphism, as identified by previous studies of the endogenous molecular basis of queen-worker development(Evans and Wheeler 1999; Barchuk et al. 2007; Foret et al. 2012). 18% (171/950) of genes with expression patterns associated with queen versus worker production were differentially expressed in nurse tissues, depending on whether the nurses were royal nurses or worker nurses.

These differentially-expressed nurse genes associated with caste development provide putative examples of genes with indirect genetic effects, which occur when genes expressed in one individual affect traits expressed by a social partner(Wolf and Moore 2010). Many of the highly-expressed and caste-associated genes we identified have protein products that have previously been found in royal jelly(Schonleben et al. 2007; Furusawa et al. 2008; Zhang et al. 2014). These nurse-produced royal jelly components are directly fed to developing larvae, providing a direct mode of action of social regulation of larval caste fate(Kamakura 2011; Huang et al. 2012). Other caste-associated nurse genes with protein products that are not known to be secreted into royal jelly may have a more circuitous effect on larval caste fate through their effect on nurse worker physiology or provisioning behavior(Haydak 1970; Brouwers et al. 1987; Hatch et al. 1999).

Quantitative genetic studies using the interacting phenotypes framework in a range of organisms, from plants to social insects to mammals, have shown that indirect genetic effects make strong contributions to heritable variation and can strongly affect evolutionary dynamics(Bleakley et al. 2010; Wolf and Moore 2010). Our study demonstrates that the interacting phenotypes framework is readily extended to consider the full transcriptional architecture and molecular basis of complex social traits, including genes with both direct and indirect effects, i.e. the “social interactome” – as opposed to only focusing on the subset of these genes that currently harbor segregating variation and contribute to observed patterns of phenotypic variation. Our results hint at a much broader contribution of nurse-expressed genes to the colony-level gene network regulating caste development than has previously been considered, consistent with the notion that caste is influenced by multiple nurse-produced and nurse-regulated factors(Linksvayer et al. 2011; Leimar et al. 2012; Buttstedt et al. 2014).

Genes with caste-associated expression had higher estimated selection coefficients than non-differentially-expressed genes and genes with expression dependent on queen presence (Fig. 4). Furthermore, among caste-associated genes, genes up-regulated in worker larvae and worker nurses had higher selection coefficients than genes up-regulated in queen larvae and royal nurses (Fig. 5). Altogether, these results indicate that both genes with direct effects and putative indirect effects on larval development – especially those associated with worker development – may have experienced positive selection and contributed to the evolution of the honey bee caste system. Our results fit with two recent honey bee studies showing that genes associated with *adult* worker traits are also rapidly evolving. The first study shows that genes encoding proteins that are more highly expressed in adult honey bee workers compared to adult queens have experience stronger selection(Harpur et al. 2014). The second study finds that the most highly-expressed genes in specialized adult tissues with derived social functions, such as the hypopharyngeal and mandibular glands, tend to be very rapidly evolving, taxonomically-restricted genes(Johnson and Tsutsui 2011; Jasper et al. 2014).

In accordance with previous transcriptomic studies(Grozinger et al. 2003; Malka et al. 2014; Manfredini et al. 2014), we also identified many genes with expression patterns dependent on queen presence, demonstrating that removal of the colony queen has broad effects on mean patterns of nurse gene expression. Most (90%, 1863/2069 genes) such queen presence-dependent expression occurred in nurse mandibular gland tissue, with a relatively small number of genes differentially expressed in larvae and other nurse tissues. At the colony level, queen removal or death results in a rapid shift from exclusively worker rearing to emergency rearing of a handful of new queens, and these broad gene expression changes may thus be associated with physiological changes associated with the production of new queens. On the longer term, if new queen production is not successful so that the colony is hopelessly queenless, some workers undergo further physiological and behavioral changes and begin laying unfertilized drone eggs(Thompson et al. 2008; Cardoen et al. 2011).

As expected, genes whose proteins make up the primary components of royal jelly, including 8 of the 9 major royal jelly proteins (MRJPs), were among the most highly-expressed genes in nurse tissues (Table S1) and were also differentially expressed (Tables 1 and 2). However, of the *mrjp* genes, only the expression of *mrjp3*, which has previously been implicated as promoting queen development(Huang et al. 2012), depended on nurse behavior: it was up-regulated in the head tissue of royal nurses (Table 1). All eight differentially-expressed *mrjp* genes, including *mrjp1*, also implicated as promoting queen development(Kamakura 2011), were differentially expressed in nurse mandibular glands or head tissue, depending on queen presence. Most were up-regulated in the queen-removed condition (Table 2), presumably related to colony-level changes associated with the rapid shift to emergency queen rearing.

Notably, 4 of the 6 described honey bee antimicrobial peptides(Evans et al. 2006) (*defensin 1, abaecin, hymenoptaecin*, and *apisimin)* were up-regulated in the HPG and / or the head tissues of nurses feeding queen-destined larvae (Table 1) and caste-associated nurse-expressed genes were enriched for Gene Ontology terms for immune function (Table S4). Interestingly, *hymenoptaecin* and another antimicrobial peptide, *apidaecin* were also upregulated in queen-destined larvae. Altogether, these results suggest that queen- and worker-destined larvae may require different levels of antimicrobial peptides, some of which may be produced by nurse workers and transferred to larvae through royal jelly(Schonleben et al. 2007; Furusawa et al. 2008; Zhang et al. 2014).

Besides broad differences in gene expression, royal nurses also had larger HPG acini and were 1.5 days younger than worker nurses (Fig. 2 and 3). Previous studies have shown that nurse HPG gland size and activity(Ohashi et al. 2000), as well as the composition of nurse glandular secretions(Haydak 1970), and patterns of nurse brain gene expression(Whitfield et al. 2006) all vary with nurse age and social environment. While it is not clear how exactly these differences are related to the observed differences in nurse provisioning behavior, individually-marked nurses did tend to specialize on feeding either queen or worker cells within a day, but not across multiple days. Thus, individual nurses may be physiologically primed to contribute to queen rearing only on the short term. Longer-term tracking of individuals during queen rearing will be necessary to definitively demonstrate the degree to which nurse specialization occurs. The key point for this study of colony-level caste regulation is that queen- vs. worker-destined larvae interact with nurses that are on average transcriptionally and physiologically distinct, resulting in distinct rearing environments and alternate caste developmental trajectories.

## Acknowledgements

Lucy Snyder, Joelle Orendain, and Brian Martinez helped with individually-marking bees and Lucy Snyder helped with behavioral observations. Tim Sheehan helped with behavioral observations and construction of the observation hives. Sandra Rehan and Nadeesha Perera measured HPG acini size and prepared tissue samples for sequencing. This research was funded in part by a University of Pennsylvania University Research Foundation grant to TAL. SV was supported by a NIH-PERT fellowship K12GM000708. AZ was funded by a NSERC Discovery grant.

## Author Contributions

TAL, SV, and KEA designed the study. SV, TAL, and KEA carried out the study. TAL, BRJ, BH, CK, and AZ analyzed the data. TAL wrote the manuscript.

## Data Accessibility

RNA Seq reads will be deposited in the NCBI SRA archive

Read counts per sample are included as a supplemental table

Raw behavioral scan data will be uploaded to Dryad.

## Supplemental Figure Captions

**Figure S1.**
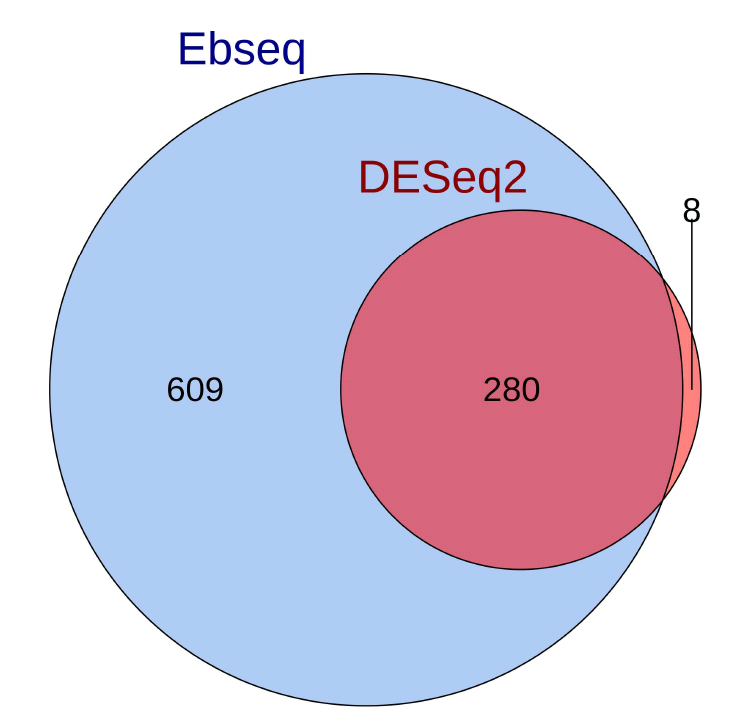
Venn diagram showing overlap of differentially-expressed genes associated with caste identified by EBSeq and DESEq2. For this comparison, DESeq2 is more conservative, identifying mainly a subset of EBSeq-identified genes.

**Figure S2.**
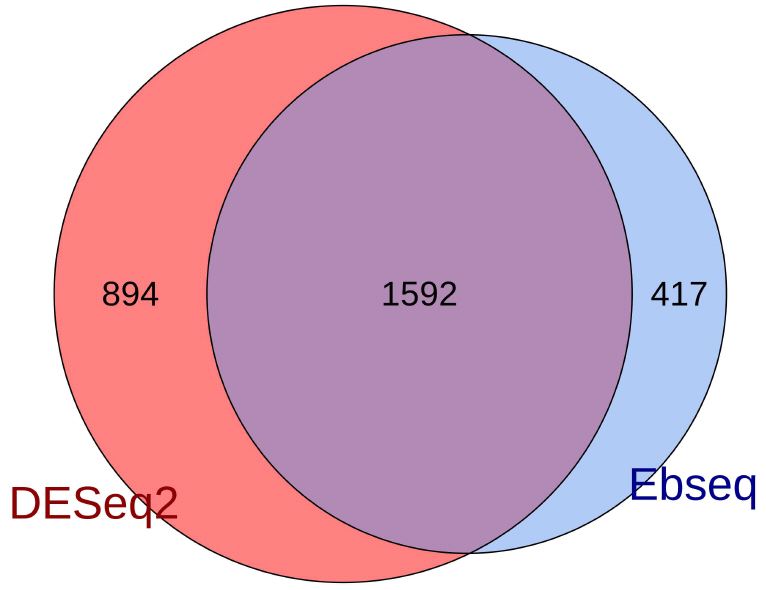
Venn diagram showing overlap of differentially-expressed genes associated with queen presence identified by EBSeq and DESEq2. For this comparison, EBSeq is somewhat more conservative than DESeq2, with less overlap than for caste-associated expression.

**Figure S3.**
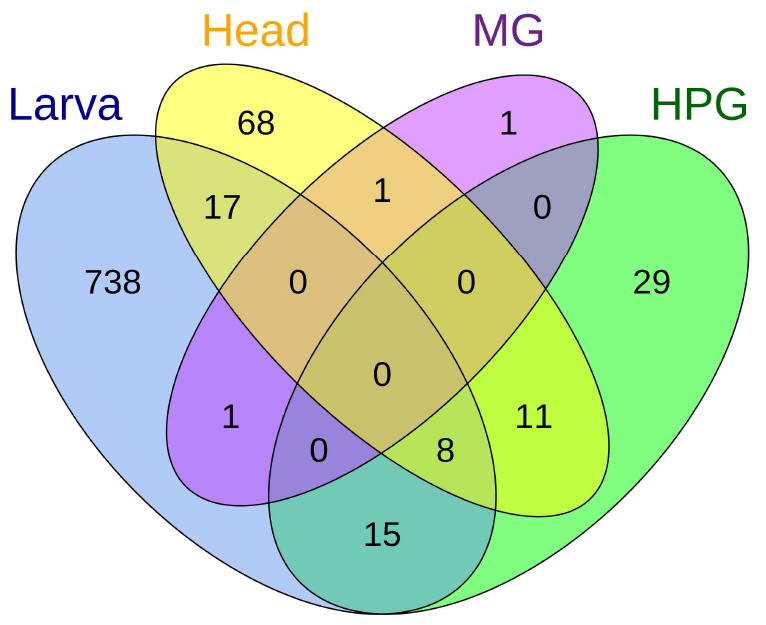
Venn diagram showing overlap for all differentially expressed genes associated with caste for each tissue. Results are based on EBSeq analysis.

**Figure S4.**
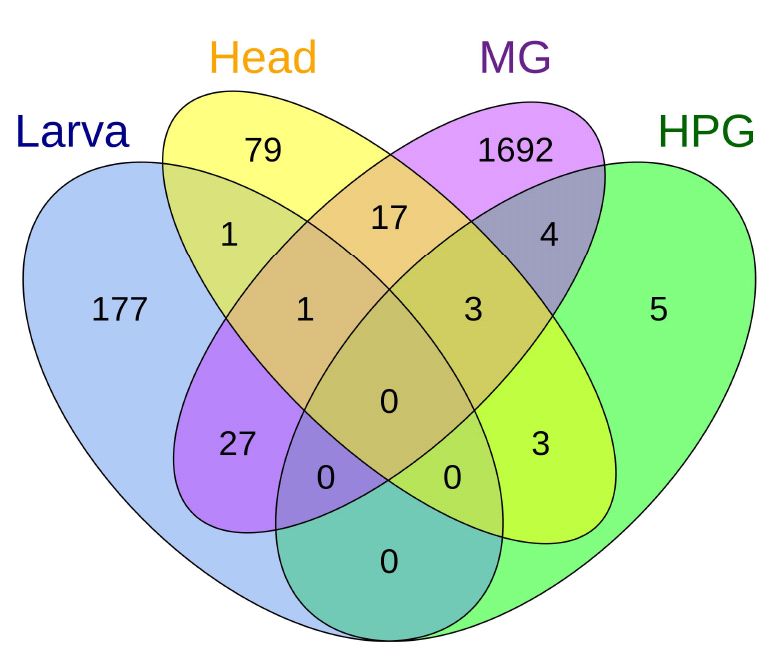
Venn diagram showing overlap for all differentially expressed genes associated with queen presence for each tissue. Results are based on EBSeq analysis.

**Figure S5.**
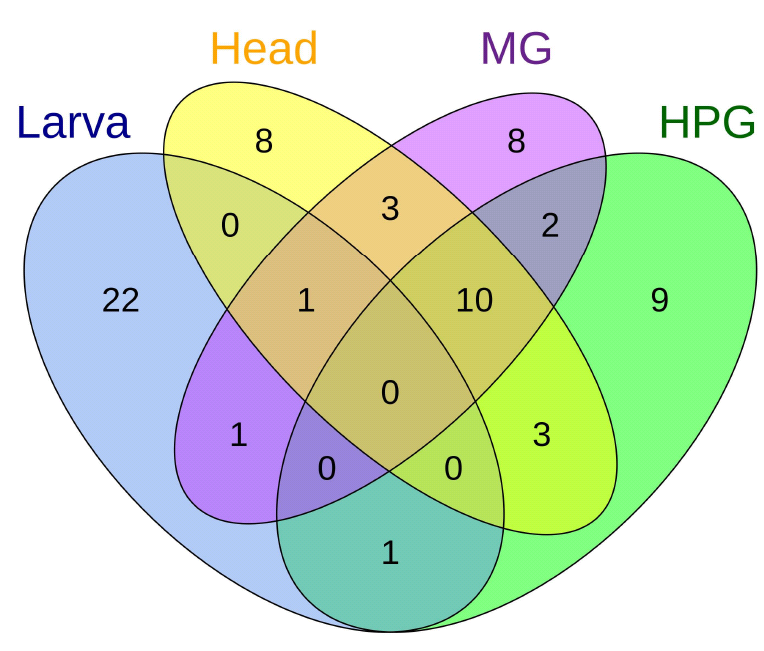
Venn diagram showing overlap of the top 25 most highly expressed genes for each tissue.

Supplemental Table 1. The top 25 most highly expressed genes by tissue (HPG = nurse hyopharyngeal gland tissue; H = remaining nurse head tissue; L = larval tissue; MG = nurse mandibular gland tissue). Mean expression level is shown as Log10FPKM. Genes whose proteins have been identified in studies of the royal jelly proteome are identified.

Supplemental Table 2. All differentially expressed genes associated with caste development, identified by EBSeq or DESeq2, grouped by tissue and sorted by expression level. Mean expression level (FPKM) is shown; log2FoldChange indicates the log2 fold change when comparing queen-associated gene expression to worker-associated gene expression; lfcSE shows the standard error for log2FoldChange; the columns DESeq2 and EBseq indicate whether the genes were identified as being differentially expressed with DESeq2 and EBseq analysis, respectively; the column “NL” indicates whether the gene was differentially expressed in nurse (N) or larval (L) tissue; “QW” indicates whether the gene was upregulated in worker larvae or worker nurse tissues (W) or queen larvae or royal nurses (Q).

Supplemental Table 3. All differentially expressed genes associated with queen presence, identified by EBSeq or DESeq2, grouped by tissue and sorted by expression level, as in Table S2.

Supplemental Table 4. GO analysis for caste-associated genes by tissue.

Supplemental Table 5. Supplemental Table 5. GO analysis for queen-presence associated genes by tissue.

